# Y and Mitochondrial Chromosomes in the Heterogeneous Stock Rat Population

**DOI:** 10.1101/2023.11.29.566473

**Authors:** Faith Okamoto, Apurva S Chitre, Thiago Missfeldt Sanches, Denghui Chen, Daniel Munro, NIDA Center for GWAS in Outbred Rats, Oksana Polesskaya, Abraham A Palmer

## Abstract

Genome-wide association studies typically evaluate the autosomes and sometimes the X Chromosome, but seldom consider the Y or mitochondrial Chromosomes. We genotyped the Y and mitochondrial chromosomes in heterogeneous stock rats (*Rattus norvegicus*), which were created in 1984 by intercrossing eight inbred strains and have subsequently been maintained as an outbred population for 100 generations. As the Y and mitochondrial Chromosomes do not recombine, we determined which founder had contributed these chromosomes for each rat, and then performed association analysis for all complex traits (n=12,055; intersection of 12,116 phenotyped and 15,042 haplotyped rats).

We found the eight founders had 8 distinct Y and 4 distinct mitochondrial Chromosomes, however only two of each were observed in our modern heterogeneous stock rat population (Generations 81-97). Despite the unusually large sample size, the p-value distribution did not deviate from expectations; there were no significant associations for behavioral, physiological, metabolome, or microbiome traits after correcting for multiple comparisons. However, both Y and mitochondrial Chromosomes were strongly associated with expression of a few genes located on those chromosomes, which provided a positive control. Our results suggest that within modern heterogeneous stock rats there are no Y and mitochondrial Chromosomes differences that strongly influence behavioral or physiological traits. These results do not address other ancestral Y and mitochondrial Chromosomes that do not appear in modern heterogeneous stock rats, nor do they address effects that may exist in other rat populations, or in other species.

**Article Summary:** Heterogeneous stock rats were created in 1984 by intercrossing eight inbred strains. This genetically and phenotypically diverse population has been used for numerous genetic studies. We developed a method (leveraging existing data) to identify the founder strain origin of Y and mitochondrial Chromosomes in modern heterogeneous stock rats. We examined effects of these chromosomes’ genotype on behavioral, physiological, and gene expression traits among 12,055 rats. We found no significant associations, except for expression of genes located on these chromosomes.

## Introduction

Heterogeneous stock (**HS**) rats (*Rattus norvegicus*) are a well-established outbred population that have been used for genome wide association studies (**GWAS**); yet, their Y and mitochondrial (**MT**) Chromosomes have been largely ignored. The Y Chromosome was poorly assembled in prior versions of the rat genome. However, the most recent rat reference genome (mRatBN7.2) dramatically improved the assembly of the Y Chromosome. In contrast, the MT Chromosome was not updated in the most recent assembly (Tutaj *et al*. 2019; de Jong *et al*. 2023).

HS rats have been outbred for almost 100 generations. They were created in 1984 by intercrossing eight inbred strains: ACI/N, BN/SsN, BUF/N, F344/N, M520/N, MR/N, WKY/N, and WN/N (Hansen and Spuhler 1984). Modern HS rat genomes are mosaics of those 8 founder haplotypes (Solberg Woods and Mott 2017), which enables precise genetic mapping of complex traits (e.g., Johannesson *et al*. 2009; Baud *et al*. 2013; Chitre *et al*. 2020). However, as Y and MT are nonrecombinant, even in a modern HS rat, they are expected to be inherited in their entirety from a single founder; the Y Chromosome from the father and the MT from the mother.

Some Y and MT haplotyping methods cannot be used in HS rats. For example, we lack complete pedigrees that have been used to trace expected Y or MT genotypes, as was done in Collaborative Cross (**CC**) mice (Broman 2022). We also lack curated lists of informative variants, as in human databases (e.g., Kloss-Brandstätter *et al*. 2011; Chen. *et al*. 2021).

In humans, Y or MT haplogroups have been tested for association with many phenotypes (e.g., Jamain *et al*. 2002, Ma *et al*. 2014, Howe *et al*. 2017, Cai *et al*. 2021, Degenhardt *et al*. 2022). Replication has proved difficult; population structure confounds such work (Hagen *et al*. 2018). For example, schizophrenia was linked to MT in a Han Chinese (Wang *et al*. 2013) cohort, but not Spanish (Mosquera-Miguel *et al*. 2012) or Swedish (Gonçalves *et al*. 2018) cohorts.

Studies in CC mice found that Y or MT genotype was not associated with sex ratio (Haines *et al*. 2021), but was associated with expression of genes located on the Y and MT Chromosomes (Keele *et al*. 2021). Mouse models designed for isolating genetic effects of Y (e.g. Martincová *et al*. 2019) and MT (e.g. Welch *et al*. 2023) found phenotypic associations, even suggesting transgenerational effects of paternal Y Chromosome genotype in daughters (Nelson *et al*. 2010). However, our review of the literature did not find comparable Y or MT analyses in outbred mice.

We identified variants that could be used to determine which founder had contributed the Y and MT to each individual HS rat. This approach is broadly similar to a prior study in DO mice (Chesler *et al*. 2016). We then tested for associations between Y and MT genotypes and a large collection of phenotypic data that have been collected over almost a decade of studies using HS rats (www.ratgenes.org). These data include behavioral, physiological, metabolome, microbiome and RNA-seq complex traits; in total, we analyzed 12,055 haplotyped and phenotyped HS rats.

## Materials and Methods

A Reagent Table is in the Supplementary Files.

### Genotype datasets

We used pre-existing whole-genome sequencing (**WGS**) data from males representing each of the 8 founder strains (∼40x coverage). SNPs and indels on the Y and MT Chromosomes were called using GATK, as previously described (Chen *et al*. 2023a). We used these data to identify polymorphic sites distinguishing the different founder Y and MT Chromosomes. We also used WGS data from 44 male and 44 female outbred HS rats (∼33x coverage); SNPs and indels were called using GATK. Short tandem repeats (**STR**s) on Y Chromosome were called in all of these samples with HipSTR and filtered with DumpSTR (Willems *et al*. 2017; Mousavi *et al*. 2020).

We also used pre-existing low-coverage (∼0.25x) data from 15,120 outbred HS rats. These used double-digest genotyping-by-sequencing (**ddGBS**; Gileta *et al*. 2020) or low-coverage WGS (**lcWGS**; Chen *et al*. 2023b) library preparation. Biallelic single nucleotide polymorphism (**SNP**) genotypes were imputed via STITCH on mRatBN7.2. We did not use the variant filters previously described. Instead, we started with all variants produced by STITCH and then used custom filters to avoid excluding variants potentially useful to distinguish founder Y or MT (see “Genotype filters”). Because Y and MT are hemizygous, heterozygous calls are unexpected (Figure S1), when observed, those genotypes were treated as missing. All procedures prior to tissue collection were approved by the relevant Institutional Animals Care and Use Committees.

### Genotype filters

Our custom filters were designed to (1) remove variants with low INFO score (for the low-coverage data), (2) remove monomorphic variants (minor allele frequency (**MAF**) = 0), (3) remove variants with a high missing rate (>25%), and (4) remove individual samples with a high missing rate (>50%). We applied all or only a subset of these filters, always in the above order, depending on the analysis. In particular, when visualizing by SNP to determine haplotypes (e.g. in alignments) we skipped the MAF filter to visualize fixed variants, and when plotting statistics (e.g. heterozygosity) by SNP in low-coverage data we skipped all but the INFO score filter. Figure S2 shows distributions of these statistics (INFO score, MAF=0, per-SNP missing rate, per-sample missing rate) for low-coverage samples, and the thresholds used. These are the filters that were used to produce haplotype groups for association analyses.

### Unrooted trees

We applied all standard filters to high-coverage genotype data. We use a matrix of Hamming distance (scale of 0 to 1) pairwise ignoring missingness, i.e., removing variants missing in either sample. We created an unrooted neighbor-joining (**NJ**) tree (Talevich *et al*. 2012). These trees were used for understanding HS founder phylogeny, but not for haplotype group-making.

### Statistical analysis

We performed a phenome-wide association study (**PheWAS**) for Y or MT haplotype via mixed linear model-based association (**MLMA**) analysis (Yang *et al*. 2014) with GCTA (Yang *et al*. 2011); see “GWAS phenotype association”. We tested normalized (“cpm” in edgeR) RNA-seq transcript abundance against Y or MT haplotype via a two-sample Wilcoxon rank-sums (i.e. Mann-Whitney) test (“wilcox.test” in R); see “Gene expression association”. We used the Benjamini & Hochberg (**BH**) false discovery rate (**FDR**) approach (“p.adjust” in R; Benjamini and Hochberg 1995). For a single, binary phenotype (“number of kidneys at birth”), we tested for association with MT haplotype using a Fisher’s exact test (“fisher.test” in R).

### GWAS phenotype association

We used a genetic relationship matrix (**GRM**) constructed (--make-grm-bin) using PLINK (Chang *et al*. 2015) to account for autosomal (--chr 1-20) relatedness (Yang *et al*. 2010), which we expected to be correlated with Y and MT haplotype due to familial structure. After filtration by missingness (--geno 0.1), violations of Hardy-Weinberg equilibrium (--hwe 1e-10; Wigginton *et al*. 2005), and MAF (--maf 0.005), 5,315,011 SNPs and 15,120 samples remained. We fit a linear model on all raw values and covariates, then inverse-normal transformed the residuals.

The traits used for the PheWAS are shown in Table S1. We encoded Y and MT haplotypes as pseudo-SNPs: reference-like haplotype (from the same haplogroup as BN) as reference allele, and alternate-like haplotype as alternate allele. We ran GCTA’s MLMA with these genotypes, the autosomal GRM, and processed phenotypes. We applied BH correction across all GWAS phenotypes, separately for Y and MT. We used FDR < 0.05 to define significance.

### Gene expression association

Our previous work mapping cis expression quantitative trait loci (**eQTL**s) showed a linear mixed model is unnecessary (Munro *et al*. 2022). Therefore, for computational simplicity, we approached gene expression analysis using methods standard in differential expression (**DE**) analysis, treating Y and MT haplotype as “conditions”, instead of eQTL mapping.

We used RNA-seq data presented as “log2” read count for all 10 tissues available from RatGTEx (Table S2), processed using the mRatBN7.2 genome build. The following filtering schema was applied (separately for Y and MT): (1) samples with a haplotype assignment were retained, (2) for each tissue, genes that had detectable expression in less than 10% of samples were excluded.

We normalized counts using Trimmed Mean of M values (**TMM**; Robinson and Oshlack 2010), then used a Mann-Whitney test for DE. This test is robust to violation of a distribution (e.g. negative binomial) in large-sample DE analysis (Li *et al*. 2022). We again used FDR < 0.05 to define significance for all genes, in all tissues, for both the Y and MT Chromosomes.

A standard eQTL expression normalization method, which involves ranking genes within a sample (Munro *et al*. 2022), is nonoptimal for highly expressed genes, ranked highly in all samples. Ranking loses raw abundance information by introducing ties between ranks. Thus, we normalized used TMM, which is a standard DE method (Corchete *et al*. 2020; Zhao *et al*. 2021).

### Data availability

Raw reads for all low-coverage samples are in the Sequence Read Archive (accession: PRJNA1022514). RNA-seq data is from RatGTex (https://ratgtex.org/download/) and archived in https://ratgtex.org/download/study-data/. An object in the UCSD Library contains all data necessary to reproduce the analysis and raw results (including unadjusted p-values) from all association tests. GWAS phenotype names are only given for significant associations to respect unpublished data collected by our numerous collaborators. HS rats are available from the NIDA Center for GWAS in Outbred Rats (https://ratgenes.org/cores/core-b/). Code to reproduce these analyses is available from GitHub.

## Results

### Two versions of Y are present in modern HS rats

All HS founders have distinct Y Chromosomes (Figure 1A). BN, ACI, and MR are relatively similar to one another, and are also similar to the reference genome (which is based on BN), while the other five founders form a separate haplogroup.

**Figure 1.**
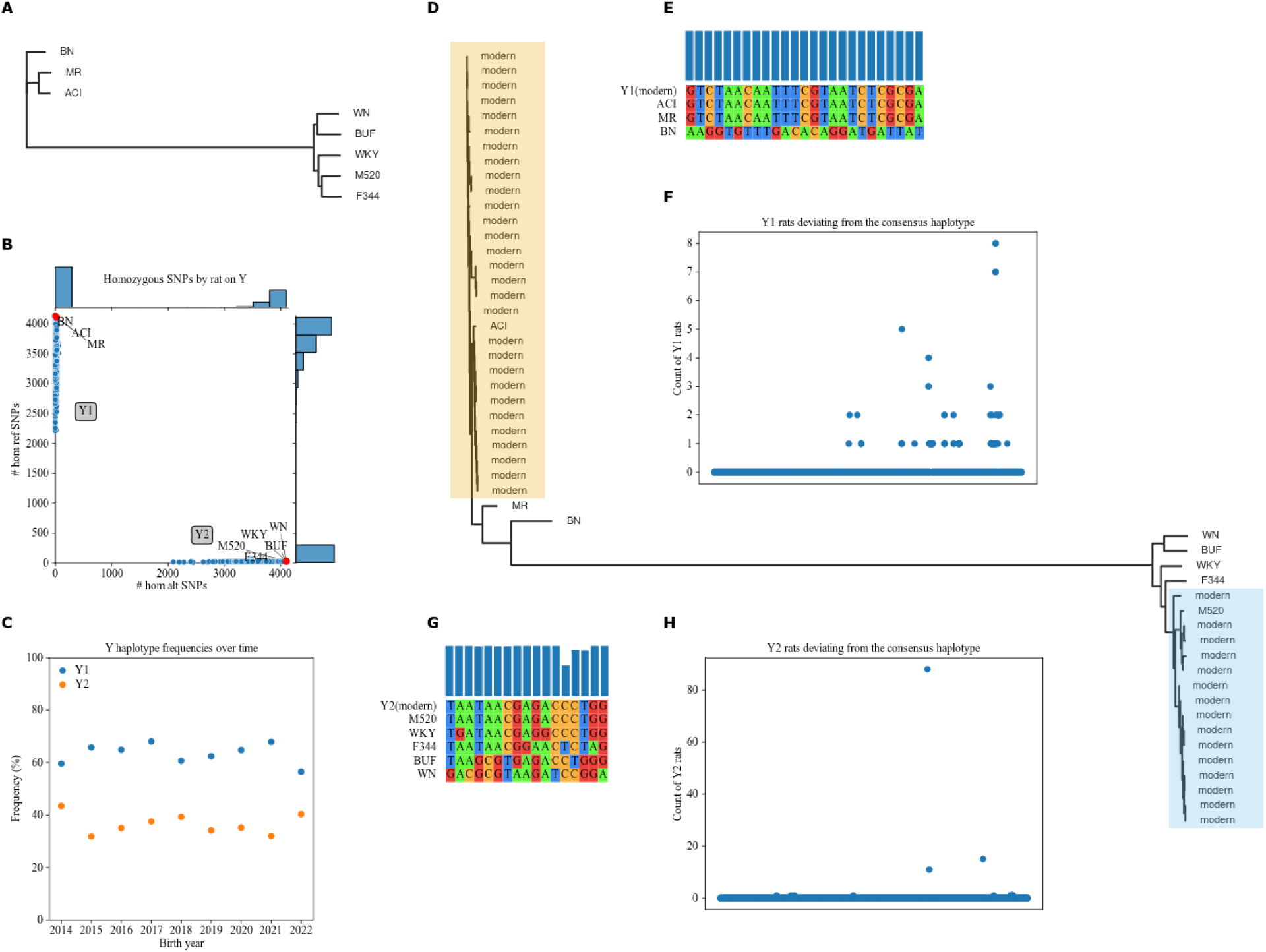
Y haplotypes present in HS founders and modern HS rats. **A.** NJ, unrooted tree using Y SNPs and indels in HS founders. Branch lengths correspond to genetic distance. **B.** Distribution of alleles by rat among Y SNPs passing filters (see Figure S2 for filters). Plot shows count of reference alleles on X-axis and count of alternate alleles on Y-axis for each rat. Side plots are histograms of allele counts among modern HS rats (blue dots). Missingness in low-coverage modern samples leads to scatter on the axes. Labeled red dots are HS founders. Y1 and Y2 haplogroups are labeled. **C.** Distribution of Y haplotypes in the HS rat population over time. Plot shows birth year on X-axis and haplotype percentage on Y-axis. **D.** NJ, unrooted tree using Y STRs in founders and 44 (29 Y1, 15 Y2) deeply sequenced modern male HS rats. Branch lengths correspond to genetic distance. Modern clades highlighted, each including a single ostensible donor founder. **E.** Pseudo-alignment within Y1 of modern and founder haplotypes, at SNPs passing filters (see Figure S2) where the Y1 founders are variable. **F.** Number of modern Y1 low-coverage genotypes deviating from the modern Y1 consensus. Plot shows SNP position along the Y chromosome on X-axis and number of low-coverage Y1 genotypes different from the haplotype on Y-axis. **G.** Pseudo-alignment within Y2 of modern and founder haplotypes, at SNPs passing filters (see Figure S2) where the Y2 founders are variable. **H.** Number of modern Y2 low-coverage genotypes deviating from the modern Y2 consensus. Plot shows SNP position along the Y chromosome on X-axis and number of low-coverage Y2 genotypes different from the haplotype on Y-axis.

We separated modern HS rats into two Y groups. We called 5,227 Y SNPs in 7,483 low-coverage samples from male modern HS rats. 4,132 SNPs and 7,471 samples remained after filtration by INFO score, MAF, and missingness (Figure S2). We grouped samples by whether they had more reference (**Y1**; 4,732 rats) or alternate (**Y2**; 2,739 rats) SNP alleles (Figure 1B). Y1 is slightly more common in the modern male HS rat population (Figure 1C).

Using STR data for a subset of 44 modern male HS rats, we found ACI to be the most recent common ancestor of modern Y1 rats, while modern Y2 rats are closest to M520 (Figure 1D).

We next found the consensus for each Y haplotype. We use the same filters on the low-coverage samples, except for skipping MAF to retain newly fixed variants. We used these data to determine the Y Chromosome haplotype for each rat. We matched these consensuses to founders in their haplogroup; the results agree with the haplotypes identified using STR data.

Y1’s modern consensus matches ACI and MR at SNPs polymorphic among the Y1 founders, BN, ACI, and MR (Figure 1E), with negligible variation across the entire chromosome (Figure 1F). Similarly, the modern Y2 consensus matches M520 (Figure 1G). Y2 has more variation; 88/2739 rats differ at one SNP (Figure 1H), possibly a mutation from the parent haplotype M520. PCA did not reveal other groupings, except by library preparation method (Figure S3A).

### Two versions of MT are present in modern HS rats

We found four MT haplotypes among the eight HS founders (Figure 2A). BUF, F344, M520, MR, and WN share mutations relative to BN, the basis of mRatBN7.2. WKY also has a distinct MT haplotype, which was not observed among modern HS rats. The ACI haplotype is barely distinct from BUF, F344, M520, MR, and WN.

**Figure 2.**
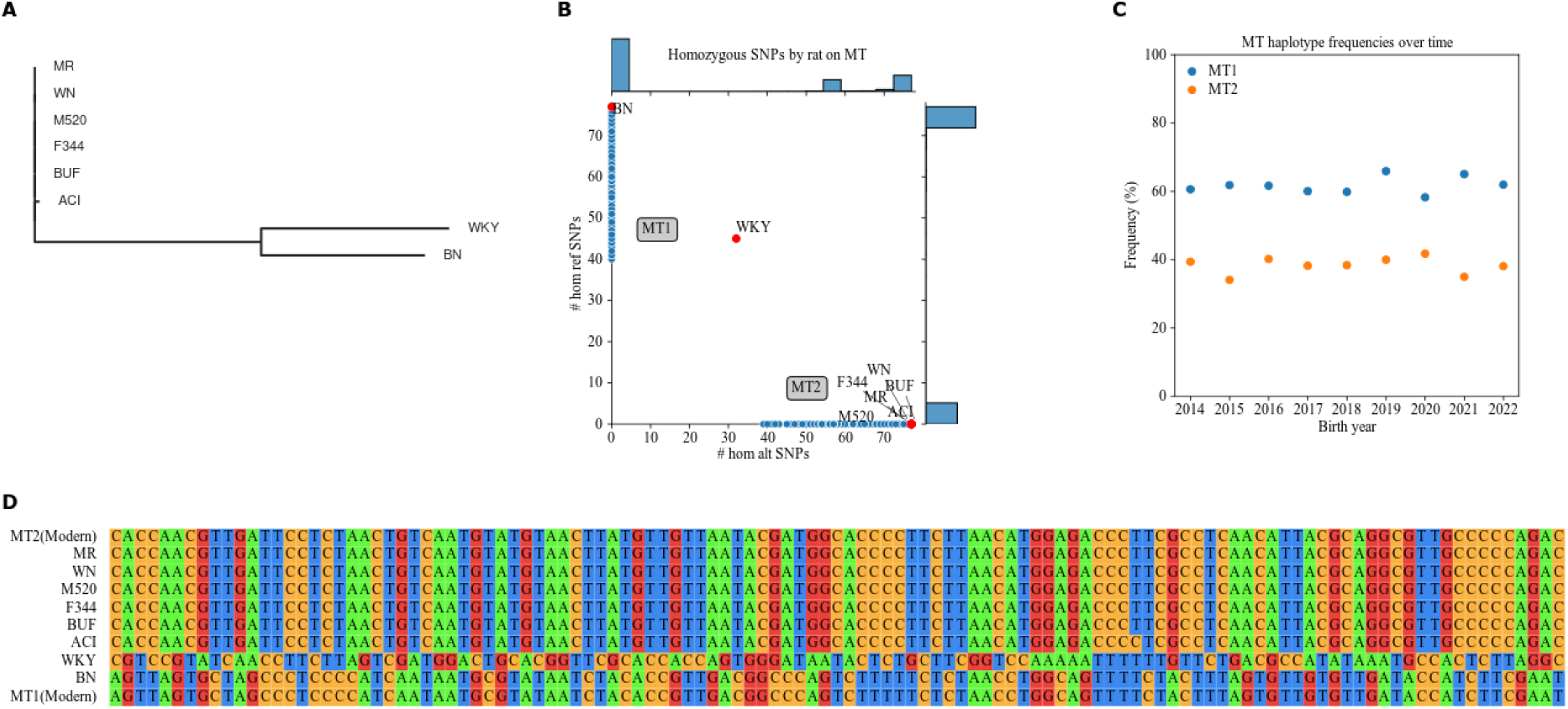
MT haplotypes present in HS founders and modern HS rats. **A.** NJ, unrooted tree using MT SNPs and indels in HS founders. Branch lengths correspond to genetic distance. **B.** Distribution of alleles by rat among MT SNPs passing filters (see Figure S2 for filters). Plot shows count of reference alleles on X-axis and count of alternate alleles on Y-axis for each rat. Side plots are histograms of allele counts among modern HS rats (blue dots). Missingness in low-coverage modern samples leads to scatter on the axes. (See Figure S4 for the bimodal distribution of MT2 missingness.) Labeled red dots are HS founders. MT1 and MT2 haplogroups are labeled. **C.** Distribution of MT haplotypes in the HS rat population over time. Plot shows birth year on X-axis and haplotype percentage on Y-axis. **D.** Pseudo-alignment of all MT SNPs called by low-coverage sequencing, colored by nucleotide. HS founders are labeled by name. The two modern haplotypes are included next to their donors.

HS founders MT phylogeny has been reported previously (Showmaker *et al*. 2020). However, their data (Ramdas *et al*. 2018) swapped WN and WKY. Our data puts WKY by itself, and WN in the large founder block with BUF, F344, M520, and MR. The Rat Genome Database (**RGD**; Vedi *et al*. 2023) Variant Visualizer (parameters: strains=HS founder strains, chromosome=MT, start=0, end=16,313) confirms the groups in Figure 2A. Complete MT genome sequencing of inbred substrains related to four of the HS founders (ACI/Eur, BN/NHsdMcwi, F344/NHsd, and WKY/NCrl) found the same relative relationships (Schlick *et al*. 2006).

We separated modern HS rats into MT groups. We called 117 MT SNPs in 15,120 low-coverage samples from modern HS rats. 77 SNPs and 14,971 samples remained after filtration by INFO score, MAF, and missingness (Figure S2). We grouped samples by whether they had more BN-like reference (**MT1**, 9,287 rats) or alternate (**MT2**, 5,684 rats) alleles (Figure 2B). MT1 is somewhat more common in the modern HS rat population (Figure 2C).

We confirmed these as the only two MT haplotypes present in the modern low-coverage SNPs genotypes. Starting with unfiltered low-coverage MT genotypes, we selected two samples with no missing SNPs, but differing genotype. All modern HS rat MT match at least one of these two. Each modern MT matches an ostensibly extant founder haplotype (Figure 2D). PCA did not reveal further groupings, however it did identify an effect of the two library preparation methods used for low coverage sequencing (Figure S3B).

### Y haplotype is associated with Y gene expression

We investigated the effect of Y haplotype on various phenotypes. Y haplotype was not significantly associated with any of the phenotypes examined (Figure 3A), except for levels of MZ531.3646417_5.08009 (Figure S5), an unannotated metabolite that was measured in the cecum. Y haplotype was associated with expression of *Ddx3y* and *Dkc1*, both of which are located on the Y Chromosome (Figure 3B-D, Table 1).

**Figure 3.**
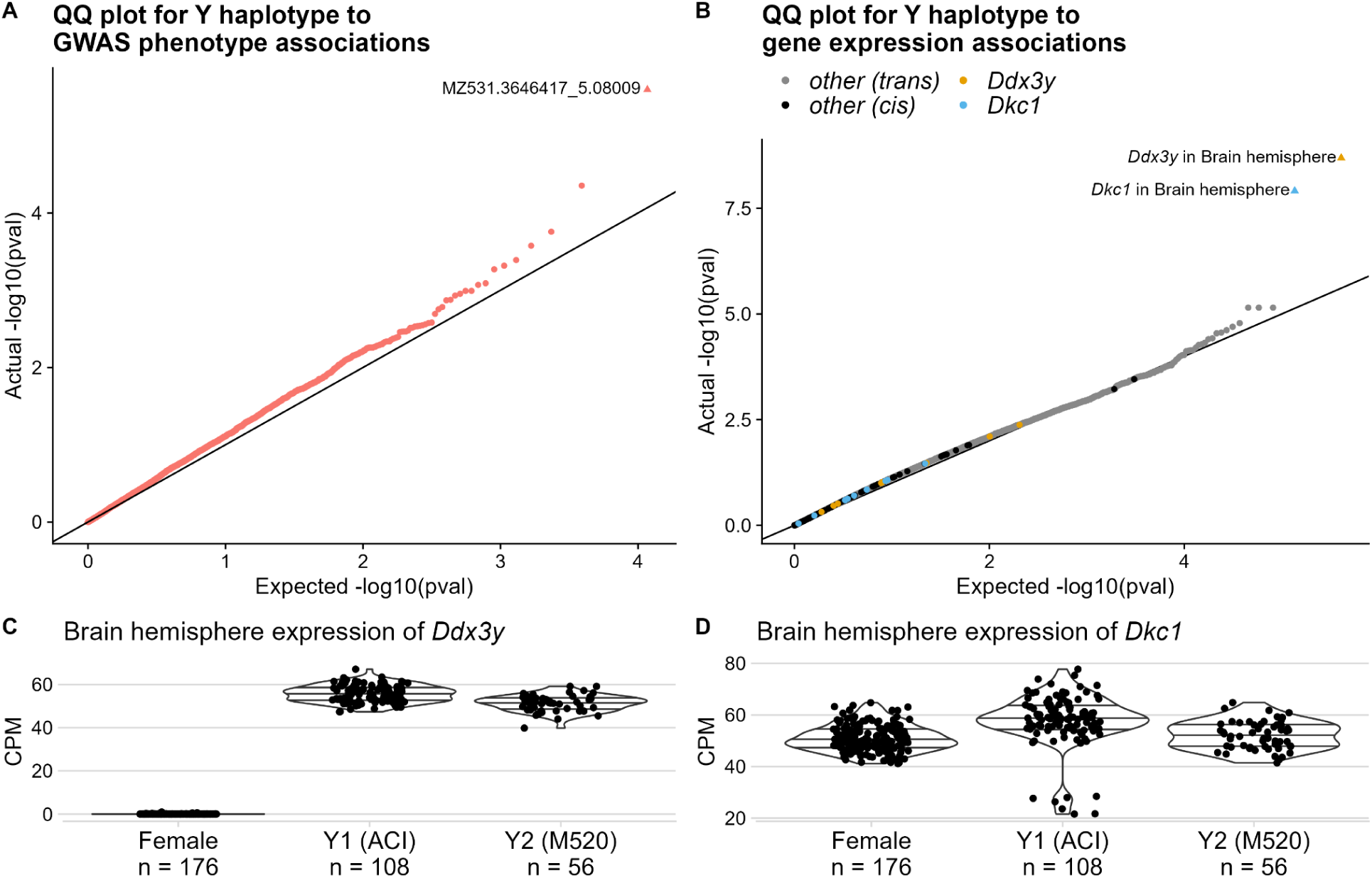
Results of Y haplotype association tests. **A.** Results of MLMAs between Y haplotype and GWAS phenotypes. Each dot represents a single trait. Plot shows actual distribution of unadjusted p-values on Y-axis, against expected distribution (null hypothesis of no association) on X-axis. Significant association (FDR < 0.05) is a triangle. **B.** Results of Mann-Whitney tests between Y haplotype and gene expression. Each dot represents a single gene in a single tissue. Plot shows actual distribution of unadjusted p-values on Y-axis, against expected distribution (null hypothesis of no association) on X-axis. Significant associations are shown as triangles. Dots for genes on Y are shown in black. Dots for the top two genes, in both the tissue with a significant association and in other tissues, are specially colored. **C-D.** *Ddx3y* and *Dkc1* CPM, split by Y haplotype, with females for context. Horizontal lines show quantiles. Plots show each sample’s normalized CPM on Y-axis; samples are split into Y haplotype groups on X-axis. Q-values are in Table 1.

**Table 1.** Genes with DE between Y haplotypes (FDR < 0.05). Information about each association is as follows: Ensembl ID (a stable identifier for the Ensembl database) of the gene, common name (from RGD) of the gene, tissue (long name and abbreviation) of the samples, chromosome the gene is on, and BH q-value of the association.

| Ensembl ID | gene | tissue | chr | q-value |
| --- | --- | --- | --- | --- |
| ENSRNOG00000057231 | *Ddx3y* | Brain hemisphere (Brain) | Y | 0.000417 |
| ENSRNOG00000055562 | *Dkc1* | Brain hemisphere (Brain) | JACYVU010000493.1 | 0.00125 |

*Ddx3y* is an RNA helicase. In humans it is involved with neuron development in males (Vakilian *et al*. 2015). Its male-specificity is sometimes used for determining sex, e.g. in humans (Hoch *et al*. 2020) and pigs (Teixeira *et al*. 2019). Consistent with this application, we found that *Ddx3y* was not expressed in female rats (Figure 3C).

Mutations in *Dkc1*’s human ortholog cause X-linked dyskeratosis congenita (Heiss *et al*. 1998); many orthologs of this gene are on the X Chromosome (Vedi *et al*. 2023). In mRatBN7.2 *Dkc1* is on an unplaced Y chromosome contig. Unlike *Ddx3y*, *Dkc1* is expressed in females (Figure 3D).

### MT haplotype is associated with MT gene expression

We investigated the effect of MT haplotype on all available phenotypes; none of the results were significant (Figure 4A). In addition, we separately tested for association with kidney number. MT1 rats have a higher rate of being born with a single kidney (see Table S3) but a one-sided Fisher’s exact test against MT haplotype was insignificant (p = 0.14). However, MT haplotype was associated with expression of several MT genes (Figure 4B-F, Figure S6, Table 2).

**Figure 4.**
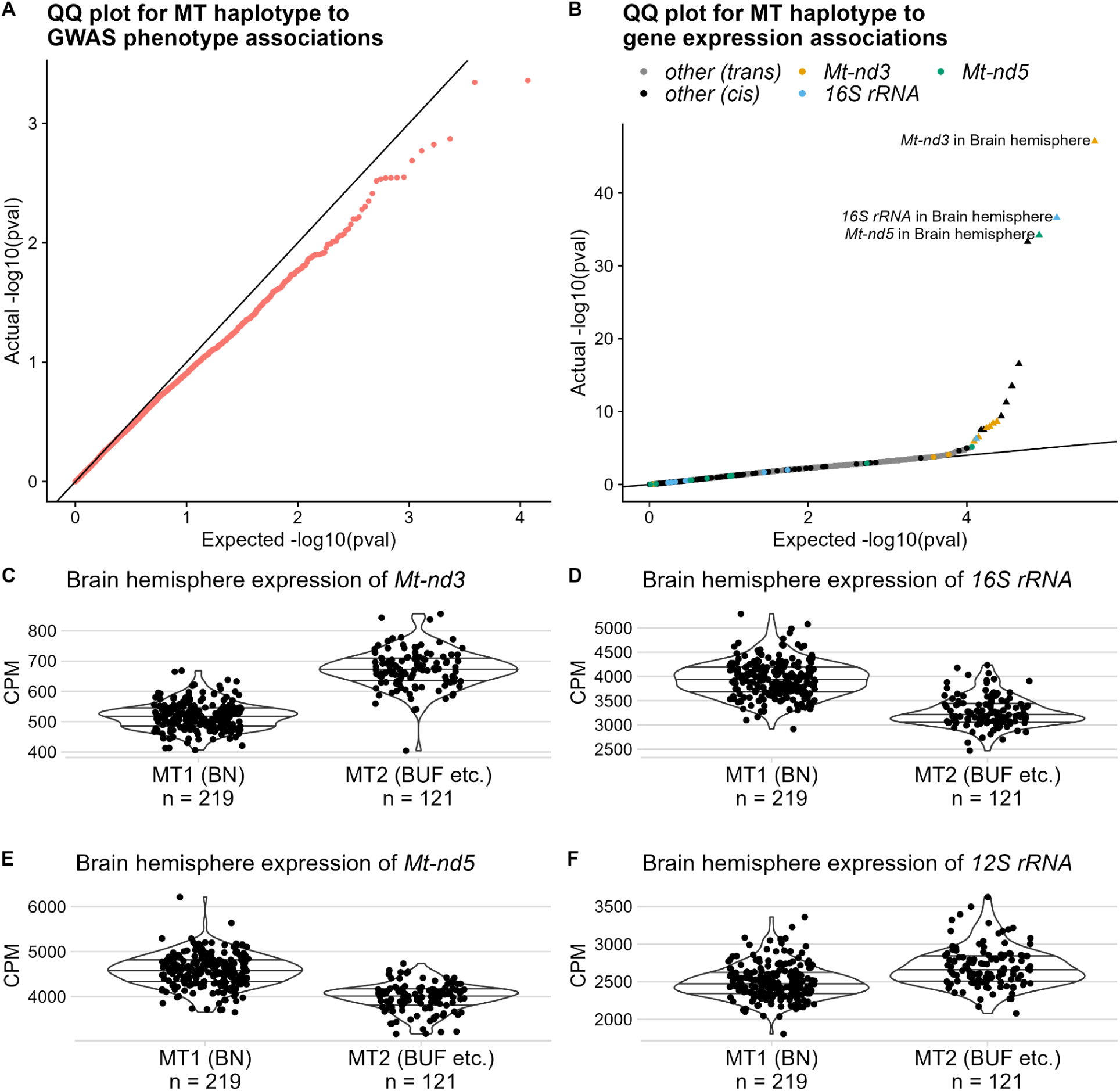
Results of MT haplotype association tests. **A.** Results of MLMAs between MT haplotype and GWAS phenotypes. Each dot represents a single trait. Plot shows actual distribution of unadjusted p-values on Y-axis, against expected distribution (null hypothesis of no association) on X-axis. **B.** Results of Mann-Whitney tests between MT haplotype and gene expression. Each dot represents a single gene in a single tissue. Plot shows actual distribution of unadjusted p-values on Y-axis, against expected distribution (null hypothesis of no association) on X-axis. Significant associations (FDR < 0.05) are shown as triangles. Dots for genes on MT are shown in black. Dots for the top three genes, in both the tissue with a significant association and in other tissues, are specially colored. **C-F.** Representative effect plots for significant associations. Plots show each sample’s normalized CPM on Y-axis; samples are split into MT haplotype groups on X-axis. Effect plots for all significant associations with MT haplotype are shown in Figure S6. Q-values are in Table 2.

**Table 2.** Genes with DE between MT haplotypes (FDR < 0.05). Information about each association is as follows: Ensembl ID (a stable identifier for the Ensembl database) of the gene, common name (from RGD) of the gene, tissue (long name and abbreviation) of the samples, chromosome the gene is on, and BH q-value of the association.

| Ensembl ID | gene | tissue | chr | q-value |
| --- | --- | --- | --- | --- |
| ENSRNOG00000033615 | *Mt-nd3* | Brain hemisphere (Brain) | MT | $1.75 \cdot 10^{-42}$ |
| ENSRNOG00000043866 | *16S rRNA* | Brain hemisphere (Brain) | MT | $2.43 \cdot 10^{-32}$ |
| ENSRNOG00000029971 | *Mt-nd5* | Brain hemisphere (Brain) | MT | $4.03 \cdot 10^{-30}$ |
| ENSRNOG00000029042 | *Mt-nd6* | Brain hemisphere (Brain) | MT | $2.51 \cdot 10^{-29}$ |
| ENSRNOG00000029707 | *Mt-nd4* | Brain hemisphere (Brain) | MT | $1.18 \cdot 10^{-12}$ |
| ENSRNOG00000030644 | *Mt-nd1* | Brain hemisphere (Brain) | MT | $1.04 \cdot 10^{-9}$ |
| ENSRNOG00000030478 | *12S rRNA* | Brain hemisphere (Brain) | MT | $1.52 \cdot 10^{-7}$ |
| ENSRNOG00000031033 | *Mt-nd2* | Brain hemisphere (Brain) | MT | $1.05 \cdot 10^5$ |
| ENSRNOG00000033615 | *Mt-nd3* | Infralimbic cortex (IL) | MT | $5.55 \cdot 10^{-5}$ |
| ENSRNOG00000033615 | *Mt-nd3* | Prelimbic cortex (PL2) | MT | $8.59 \cdot 10^{-5}$ |
| ENSRNOG00000033615 | *Mt-nd3* | Orbitofrontal cortex (OFC) | MT | 0.000210 |
| ENSRNOG00000033615 | *Mt-nd3* | Prelimbic cortex (PL) | MT | 0.000318 |
| ENSRNOG00000029042 | *Mt-nd6* | Lateral habenula (LHb) | MT | 0.000477 |
| ENSRNOG00000031053 | *Mt-nd4l* | Brain hemisphere (Brain) | MT | 0.000477 |
| ENSRNOG00000033615 | *Mt-nd3* | Lateral habenula (LHb) | MT | 0.00438 |
| ENSRNOG00000043866 | *16S rRNA* | Nucleus accumbens core (NAcc2) | MT | 0.00679 |
| ENSRNOG00000033615 | *Mt-nd3* | Nucleus accumbens core (NAcc2) | MT | 0.0146 |

Complex I is the first enzyme in the electron transport chain. In 7 of 10 tissues tested, its *Mt-nd3* subunit is up-regulated in MT2 relative to MT1. Every other MT-encoded subunit (*Mt-nd1*, *Mt-nd2*, *Mt-nd4*, *Mt-nd4l*, *Mt-nd5*, *Mt-nd6*) is down-regulated in MT2. The MT haplotypes have different subunit ratios. Also, both MT-encoded ribosomal RNAs (**rRNA**s) have significant DE.

## Discussion

We performed a large-scale study to identify phenotypes influenced by the nonrecombinant Y and MT chromosomes in 12,055 HS rats. One of our major findings was that the 8 founders of the HS population had two major Y Chromosome and three major MT Chromosome haplotype groups. In modern HS rats, we observed two Y haplogroups, with the Y1 group most closely matching ACI and the Y2 group most closely matching M520 (Figure 1). Similarly, in modern HS rats we observed two MT Chromosomes. The MT1 haplotype was most similar to BN and the MT2 haplotype matched 4 of the founders (MR, WN, M520 and F344) which could not be distinguished by any of the SNPs, indels or TRs that we examined (Figure 2).

We assigned 12,055 phenotyped and genotyped rats to Y1 or Y2 (for males) and MT1 and MT2 haplotypes and then sought to identify associations with an array of behavioral and physiological phenotypes. Remarkably, there were virtually no significant associations (Figure 3, 4). Notably, we did not find evidence to support an earlier publication by Showmaker et al. (2020) which suggested that the MT1 haplotype (BN derived) influences the chance that a rat is born with only one kidney (Table S3). We also considered gene expression, which allowed us to investigate the expression of genes located on the MT and Y Chromosomes. This analysis identified several genes located on these Chromosomes that were significantly differentially expressed.

For the Y Chromosome we identified DE of *Ddx3y*, which is involved in male human neuronal development (Vakilian *et al*. 2015), and *Dkc1* (Figure 3). For the MT Chromosome, we identified subunits of the critical respiratory enzyme Complex I, as well as rRNA, which are possible artifacts of imperfect poly-A tail selection (Figure 4, Figure S6). While these eQTLs do not appear to cause detectable changes in the behavioral and physiological traits that we studied, they provide an important positive control, demonstrating that we can accurately call Y and MT haplotypes. Overall, our results show that previous genetics studies in HS rats which did not examine the Y and MT Chromosomes, did not in fact overlook important genetic effects.

A strength of our study is the fact that the genetic structure of HS rats makes them well suited for studying Y and MT. Whereas human studies can be confounded by correlations between MT and nuclear genotype (Hagen *et al*. 2018), the HS breeding strategy (Solberg Woods and Mott 2017) and our use of MLMA for PheWAS avoided these problems. In addition, all of the observed Y or MT haplotypes are common (Figure 1C, Figure 2C), unlike the situation in DO mice (Chesler *et al*. 2016) or humans (Howe *et al*. 2017), providing better power to detect associations in HS rats.

Our results indicate that only a few of the founder Y and MT Chromosomes have persisted into modern HS rats. This could reflect genetic drift or inadvertent selection due to differences in fitness or fecundity; our data can not distinguish between these two possibilities. Thus, it is possible that some of the unobserved Y and MT Chromosomes would have shown phenotypic consequences had they been present among the modern HS rats that we studied.

In summary, we describe Y and MT haplotype structure in modern HS rats, and present results from well-powered association analyses with various phenotypes. Haplotypes are inherited from specific HS founders and cause differential expression of several genes of biological importance, including Complex I subunits and genes with orthologs to human sex-linked disorders. Methods described here may be extended to other rat populations for further investigation of Y and MT.

## Supporting information

Reagent Table

## Acknowledgements

The authors would like to thank Robert Vogel for advice in statistical analysis, Gregory Keele and Gary Churchill for useful insights, and Ryan Eveloff for providing unpublished STR data. We used the Triton Shared Computing Cluster (https://doi.org/10.57873/T34W2R) from the San Diego Supercomputer Center to run GCTA. This work was supported by P50DA037844.

