## Supplementary Figures/Tables for "Y and Mitochondrial Chromosomes in the Heterogeneous Stock Rat Population"

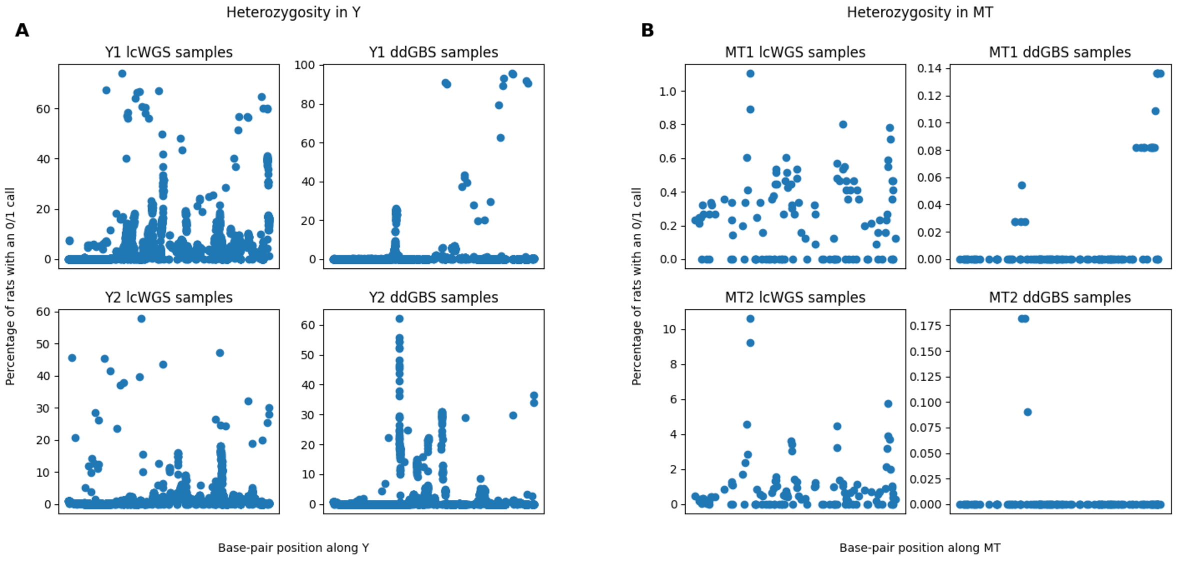


**Figure S1.** Heterozygous calls in low-coverage Y and MT chromosomes are meaningless. **A-B.** Frequency of heterozygosity, split by haplogroup and library preparation method. Plot shows SNP position along the chromosome on X-axis and percentage of rats with an 0/1 genotype call on Y-axis. Note that SNPs with heterozygosity are more dependent on sequencing method than haplogroup; if heterozygosity distinguished a true subgroup, we would expect a focused set of heterozygous calls across both sequencing methods for a single haplotype.

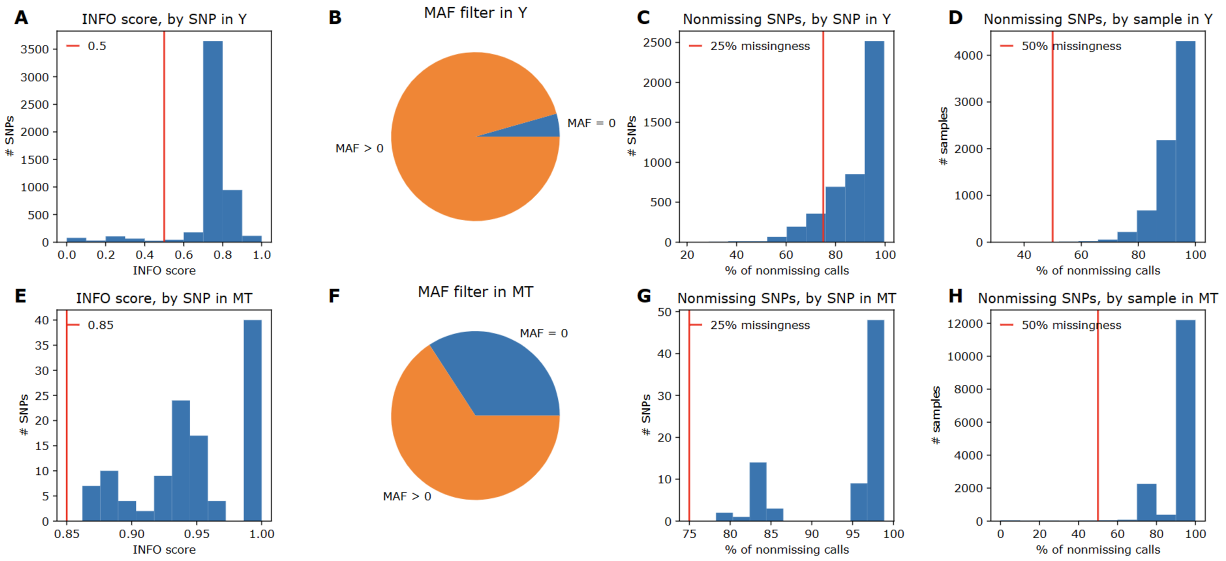


**Figure S2.** Filtration of raw low-coverage imputation output. Filters were applied from left to right; e.g. SNPs filtered out for low INFO score were not included in MAF filtration, and SNPs filtered out for by-SNP missingness were not included in calculations of by-rat missingness. Vertical lines correspond to thresholds used. **A.** A filter of INFO score >= 0.85 (threshold chosen to allow all MT SNPs) removed no MT SNPs. **B.** A filter of MAF >0 removed 40 MT SNPs **C.** A filter of by-SNP missingness <= 25% removed no MT SNPs. **D.** A filter of by-rat missingness <= 50% removed 149 MT rats. **E.** A filter of INFO score >= 0.5 (threshold chosen to be past the peak of good SNPs) removed 300 Y SNPs. **F.** A filter of MAF >0 removed 217 Y SNPs **G.** A filter of by-SNP missingness <= 25% removed 578 Y SNPs. **H.** A filter of by-rat missingness <= 50% removed 12 Y rats.

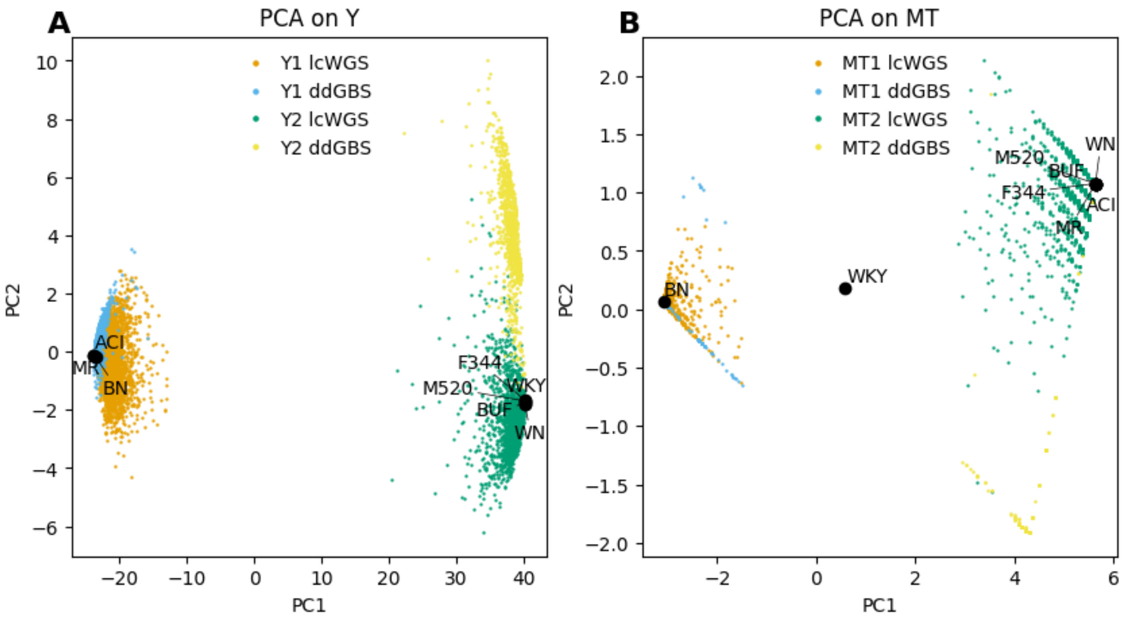


**Figure S3.** Biplot of PCA results for both chromosomes. SNPs in low-coverage genotypes were encoded as 0 (reference) or 1 (alternate), with mean imputation used for missing values. Plot shows PC1 on X-axis and PC2 on Y-axis. Modern rats colored by haplotype and library preparation method. HS founders are large, labeled black dots, projected on the same PCs.

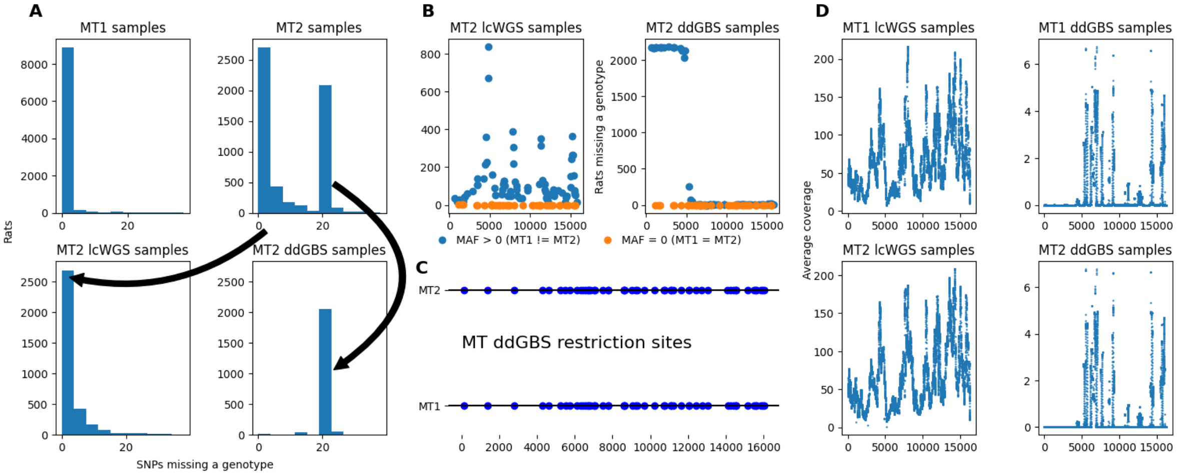


**Figure S4.** Patterns of MT missingness. Distribution of MT2 missingness is bimodal, due to a grouping of SNPs missing in ddGBS-samples. This block of SNPs is differentially imputed; both MT haplotypes lack the necessary distribution of restriction sites, and thus have essentially no coverage. However, the reference MT1, and all SNPs with MAF=0 between MT1 and MT2, were imputed. **A.** Histograms of per-rat missingness for samples with the MT1 and MT2 haplotypes, and for MT2 samples sequenced by the lcWGS and ddGBS library preparation methods. **B.** Missingness across the MT chromosome for samples with the MT2 haplotype, shown for the lcWGS and ddGBS library preparation methods. Plot shows SNP position along the MT chromosome on X-axis and count of MT2 rats missing a genotype on Y-axis. Orange dots are SNPs identical between MT1 and MT2, while blue dots show SNPs which vary between haplotypes. **C.** Locations of ddGBS restriction sites. AY172581.1 is used for MT1. MT2 is mutated based on MT SNP and indel variants in founders. X-axis is SNP position along the MT chromosome. **D**. Average coverage across MT, split by haplotype and sequencing method. The plots show base-pair position of each locus in the reference AY172581.1 on X-axis and average read coverage for each sample's library on Y-axis.

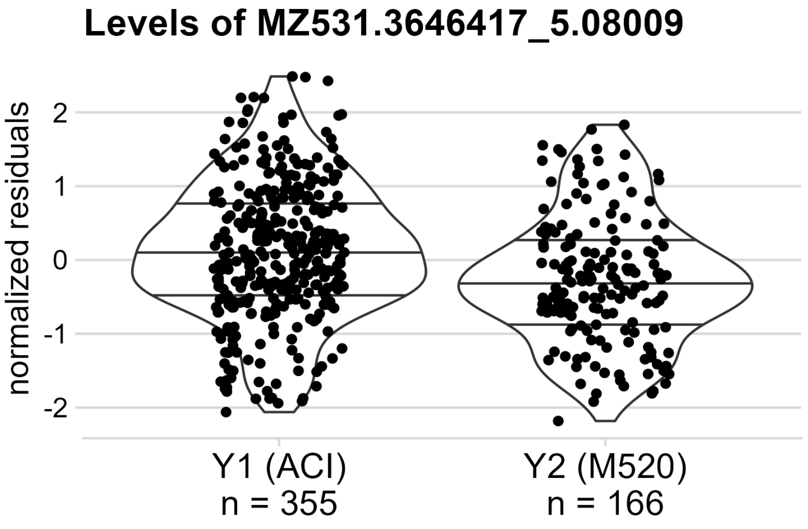


**Figure S5.** Levels of an unannotated metabolite associated with Y haplotype. Covariates were regressed out as described in "GWAS phenotype association" from Methods. Quantile lines included. Plot shows each sample's normalized residuals on Y-axis; samples are split into Y haplotype groups on X-axis. Q-value is 0.015.

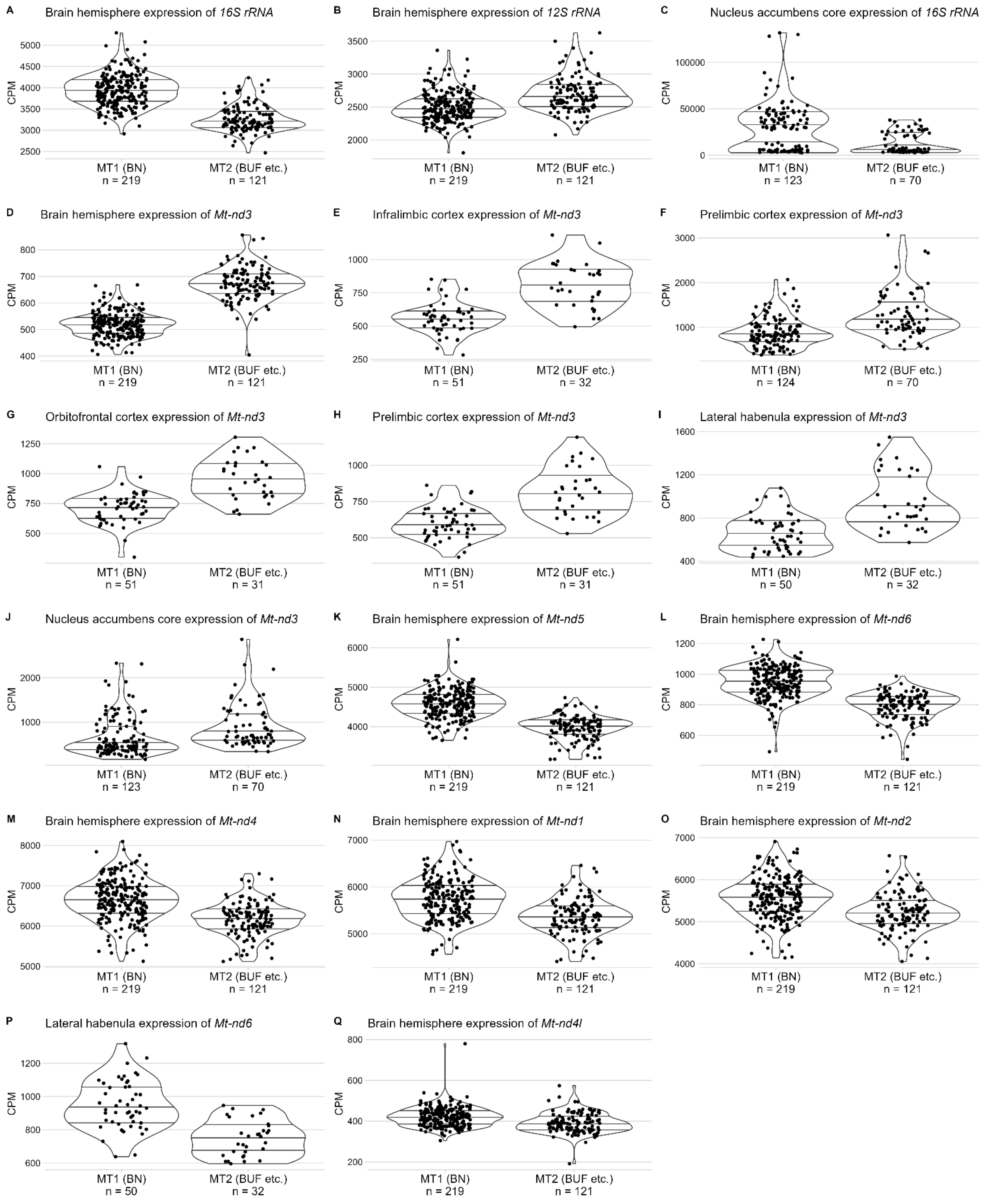


**Figure S6.** Effect plots for all significant associations between gene expression and MT haplotype. Quantile lines included. Plots show each sample's normalized CPM on Y-axis; samples are split into MT haplotype groups on X-axis. P-values of associations are in Table 1. **A-C.** MT-rRNA expression is affected by MT haplotype. **D-J.** *Mt-nd3* is upregulated in MT2 samples **K-Q.** Other Complex I subunits are downregulated in MT2 samples.

| project | traits | Y1 | Y2 | MT1 | MT2 |
|---|---|---|---|---|---|
| Association Between Behavioral Regulation and Cocaine Cue Preference | 426 | 842 | 462 | 1598 | 919 |
| Food and Water Consumption in Heterogeneous Stock Rats | 8 | 286 | 125 | 509 | 308 |
| Genetic Basis of Cecum Metabolome Composition in Heterogeneous Stock Rats | 4861 | 355 | 166 | 688 | 389 |
| Genetic Basis of Cecum Microbiome Composition in Heterogeneous Stock Rats | 108 | 1292 | 641 | 2399 | 1458 |
| Genetic Studies of Incentive Salience | 108 | 532 | 263 | 965 | 610 |
| Genetics of Adiposity Related Traits in Heterogeneous Stock Rats | 11* | 2246 | 1128 | 4115 | 2510 |
| Genetics Underlying Individual Differences in Skeletal Muscle | 4 | 1183 | 598 | 2167 | 1315 |
| Genomic Analysis of Avoidance Learning in Addiction | 9** | 244 | 143 | 671 | 408 |
| Identification of Genes Regulating Bone Matrix Composition and Quality | 29 | 262 | 178 | 537 | 343 |
| Identification of Genetic Features of Delay Discounting Using a Heterogeneous Stock Rat Model | 60 | 610 | 290 | 1106 | 632 |
| Identification of Genetic Variants thats Contribute to Compulsive Cocaine Intake in Rats | 48 | 242 | 159 | 464 | 318 |
| Neurogenetic Substrates of Cocaine Addiction | 44 | 140 | 90 | 275 | 180 |
| Socially-Acquired Nicotine Self-Administration | 11 | 598 | 299 | 1088 | 660 |
| The Genetic Basis of Opioid Dependence Vulnerability in a Rodent Model | 28 | 260 | 190 | 525 | 338 |
| Use of Next-Gen Sequencing to Identify Genetic Variants that Influence Compulsive Oxycodone Intake in Outbred Rats | 95 | 169 | 100 | 335 | 192 |

**Table S1.** Summary of GWAS phenotypes, by original project. We used all traits for which genotyped rats were available**. Some rats had phenotypes collected for more than one project. Not all rats have a MT haplotype assignment, and not all male rats have a Y haplotype assignment. Such rats were excluded from analysis. Information about each project is as follows: title, number of traits included in association tests, and number of rats phenotyped by the project who have the Y1, Y2, MT1, and MT2 haplotypes, respectively. Some phenotypes were measured in less than the total number of rats, e.g. due to early death.
* One trait in "Genetics of Adiposity Related Traits in Heterogeneous Stock Rats" was only defined in females, and thus was not included in Y tests.
** Six traits from "Genomic Analysis of Avoidance Learning in Addiction" were originally analyzed both among all rats and sex-specifically. We only used sex-combined traits in our analysis. This removed twelve sex-specific traits, leaving the nine analyzed here.

| tissue | n Y1 | n Y2 | n Y tests | n MT1 | n MT2 | n MT tests |
|---|---|---|---|---|---|---|
| Basolateral amygdala (BLA) | 60 | 37 | 19858 | 123 | 68 | 19808 |
| Brain hemisphere (Brain) | 108 | 56 | 20386 | 219 | 121 | 20437 |
| Eye (Eye) | 16 | 8 | 18539 | 35 | 17 | 18523 |
| Infralimbic cortex (IL) | 29 | 14 | 20827 | 51 | 32 | 20822 |
| Lateral habenula (LHb) | 28 | 14 | 20902 | 50 | 32 | 20957 |
| Nucleus accumbens core (NAcc) | 25 | 13 | 20704 | 45 | 32 | 20640 |
| Nucleus accumbens core (NAcc2) | 59 | 38 | 21000 | 123 | 70 | 20866 |
| Orbitofrontal cortex (OFC) | 29 | 14 | 20650 | 51 | 31 | 20642 |
| Prelimbic cortex (PL) | 27 | 15 | 20728 | 51 | 31 | 20772 |
| Prelimbic cortex (PL2) | 60 | 37 | 19905 | 124 | 70 | 19862 |

**Table S2.** Summary of gene expression DE tests, by tissue. Tissue datasets from RatGTex. Shown is number of rats with each haplotype and number of genes where >= 10% of haplotyped samples have expression, i.e., the number of samples in each DE group and the number of DE tests conducted. Not all rats have a MT haplotype assignment, and not all male rats have a Y haplotype assignment. Such rats were excluded from analysis. Some tissues were used in multiple projects; their data is split by project, e.g. prelimbic cortex is in groups of PL1 and PL2.

| | 1 kidney | 2 kidneys |
|---|---|---|
| MT1 (BN) | 73 | 3365 |
| MT2 (nearly ACI) | 35 | 2058 |

**Table S3.** Contingency table of number of kidneys at birth and MT haplotype, for all rats where both are known. The one-sided p-value for a Fisher's exact test is 0.14 for association between MT haplotype and number of kidneys.
